# The ventral stream receives spatial information from the dorsal stream during configural face processing

**DOI:** 10.1101/222851

**Authors:** Valentinos Zachariou, Nicole Mlynaryk, Marine Vernet, Leslie G. Ungerleider

**Affiliations:** Section on Neurocircuitry, Laboratory of Brain and Cognition, National Institute of Mental Health, National Institutes of Health, Bethesda, MD 20892 USA

## Abstract

Configural face processing is considered to be vital for face perception. If configural face processing requires an evaluation of spatial information, might this process involve interactions between ventral stream face-processing regions and dorsal stream visuospatial-processing regions? We explored this possibility using thetaburst stimulation (TBS) with fMRI in humans. Participants were shown two faces that differed in either the shape (featural differences) or the spatial configuration (configural differences) of their features. TBS applied on dorsal location-processing regions: 1) reduced fMRI activity within ventral stream face-processing regions during configural but not featural face processing; and 2) reduced functional connectivity between these face regions significantly more for configural than featural face processing. No changes occurred when TBS was delivered on the vertex control site for either face task. We conclude that ventral stream face-processing regions receive visuospatial information from dorsal stream location-processing regions during configural face processing.

**Significance statement:** Face perception is thought to be mediated exclusively by neural substrates within the ventral visual pathway. However, by using non-invasive brain stimulation (thetabust transcranial magnetic stimulation) in healthy human adults, we demonstrate that the face-processing regions of the ventral visual pathway receive information from visuospatial-processing regions of the dorsal visual pathway during configural face processing, a vital function in face perception. Our findings thus indicate that veridical face perception may depend on both the ventral and dorsal visual pathways.

## Introduction

Evidence from lesion, neurophysiological and imaging studies indicates that face perception is mediated by specialized neural machinery within the primate visual system (Kanwisher et al, 1997; Haxby et al. 2000; Rossion, et al. 2003; Jonas et al. 2016; Barton 2008; Tsao et al. 2008). For example, in humans, the occipital face area (OFA; Gauthier et al. 2000) within the inferior occipital gyrus, the fusiform face area (FFA; Kanwisher et al, 1997) within the lateral fusiform gyrus, and the anterior inferior temporal face region (aIT; Sergent et al. 1992; Haxby et al. 2000; Tsao et al. 2008) within the rostral ventral temporal cortex are all activated more strongly by faces than by non-face categories of objects. Importantly, damage to these regions can result in prosopagnosia, a face-specific perceptual impairment (Della Sala and Young 2003; Duchaine et al. 2006).

Exactly how these face-specific brain regions mediate face perception is not well understood. One theory suggests that these neural substrates mediate at least two cognitive processes: featural processing, which entails perceiving the features of a face, such as the shape of the eyes, nose and mouth, and configural processing, which entails perceiving the spatial arrangement of the facial features, such as the distance between the eyes, nose and mouth (Maurer et al. 2002; Yovel and Kanwisher 2004; Maurer et al. 2007; Zhang, et al. 2015).

The processing of shape, irrespective of the object category, is considered to be a primary function of areas within the ventral occipitotemporal cortex (VOTC) that are part of the ventral visual pathway (Ungerleider and Mishkin 1982). As such, it is not surprising that face-selective areas within this pathway process the shape features of a face. In contrast, configural face processing is more of a puzzle, as it likely depends on visuospatial processing, which is not considered to be a primary function of the ventral visual pathway. Indeed, representations of space within VOTC are rudimentary and imprecise (Sereno and Lehky 2010). Is it therefore possible that the spatial information required for configural face processing is mediated by brain regions outside VOTC that are associated with visuospatial perception, such as areas within posterior parietal cortex (PPC) that are part of the dorsal visual pathway (Ungerleider and Mishkin, 1982; Mishkin et al. 1983). Neurons within this pathway encode and reconstruct stimulus locations with a high degree of spatial precision (Sereno and Lehky 2010). Therefore, it may be that these high-precision spatial mechanisms within the dorsal visual pathway contribute to configural face processing.

We previously explored this possibility in healthy human adults using fMRI and on-line repetitive transcranial magnetic stimulation (rTMS; Zachariou et al., 2016). Our findings indicated that, within localized, spatial-processing regions of PPC, configural face difference detections led to significantly stronger activation compared to featural face difference detections, and the magnitude of this activation predicted the participants’ behavioral performance. Importantly, rTMS centered on these PPC regions impaired performance on configural but not featural face difference detections. We concluded that spatial mechanisms within the dorsal visual pathway contribute to the configural processing of facial features. However, exactly how dorsal spatial information is integrated with other face-related information remains to be determined. One hypothesis, which we previously proposed (Zachariou et al. 2016), is that the integration of spatial and other face-related information is a consequence of information exchange between the spatial-processing regions in PPC and face-processing regions in VOTC. Accordingly, in this study, we provide strong evidence to support this hypothesis by demonstrating that interference induced by thetaburst magnetic stimulation (TBS) on localized location-processing regions of the dorsal visual pathway significantly reduced both the magnitude of fMRI activity and degree of functional connectivity of face-processing regions within the ventral visual pathway, specifically for configural but not featural face difference detections. Consequently, face-processing regions within the ventral visual pathway appear to receive visuospatial information from the dorsal visual pathway during configural face processing.

## Materials and Methods

### Participants

Twenty-four healthy adults were recruited for the experiment (12 females, age range 22-42 years). Four participants failed training and were excluded. One participant asked to withdraw from the study due to health issues unrelated to the study and two participants failed to complete all TBS sessions before moving away from the area. Thus, seventeen healthy adults (7 females, age range 22-42 years) participated in the experiment. All were right-handed and had normal vision (corrected, if necessary) and all gave informed consent under a protocol approved by the Institutional Review Board of the National Institute of Mental Health.

### Procedure

The experiment, implemented using E-prime 2.0, was run on a Windows-7 based PC. Stimuli were presented via an analog projector on a 450 x 344 mm screen (29° visual angle horizontally by 18° vertically at a distance of 128 cm away from the participants’ eyes), situated at the bore opening of the MRI scanner at a resolution of 1024 x 768 pixels (0.03° per pixel). Participants viewed the projection screen through a mirror attached to the head coil of the MRI scanner.

The experiment consisted of five sessions, which occurred on different days. In the first session, participants were trained on the configural/featural face difference detection tasks (described in detail below) using a training procedure similar to that described in Zachariou et al. (2016). Following training, participants completed a localizer fMRI session, identical to the one described in Zachariou et al. (2016). Then, following the localizer session, participants completed three TBS sessions on different days.

### Localizer session

During the localizer session, participants completed four functional runs, two runs each for localization of the dorsal and ventral stream ROIs. Each run of both localizer tasks consisted of 14 blocks of trials (10 trials per block) in counterbalanced order. Participants were not required to maintain fixation during these trials in order to follow the same experimental procedure as that of the TBS sessions, described below. In both localizer tasks, participants viewed two images presented simultaneously on either side of the screen center for 1.7 s. In half the trials of a block, the images appeared in a top-left, bottom-right configuration and in the remaining half, in a top-right, bottom-left configuration. Each block of trials lasted 22 s and was directly preceded and followed by 8 s of fixation. Trials were separated by 300 msec of fixation.

### Dorsal stream localizer

For the dorsal stream localizer, we compared activations evoked by a same-different distance detection task with activations evoked by a same-different brightness detection task (adapted from Haxby et al. 1991 and previously used in Zachariou et al. 2015; 2016). The two tasks were presented in separate, counterbalanced blocks and were visually very similar: the display consisted of two panels, each containing a dot and a vertical black line. The panels always depicted the dot at opposite horizontal and vertical positions. On each trial, the distance between the dots and lines was randomly drawn from a uniform distribution between 18 and 80 pixels. In addition, the brightness of the dots, on each trial, was randomly chosen to be one of eight brightness levels. Thus, the dot-line distance and dot brightness differed across trials for both tasks. Panels in identical trials in both distance and brightness tasks had the same dot-line distance and the same dot brightness. On each trial, participants compared the two panels and indicated with a button press if the panels differed in distance or brightness.

In same-different distance detection blocks, the brightness of the dot was always identical across the two panels within a trial (but varied across trials), and participants compared the horizontal distance between the dot and the vertical line across the two panels. This horizontal distance differed in half of the trials of each block, and participants indicated detection of this difference by a button press (responses withheld on identical distance trials). A distance difference across panels was created by adding 18-30 pixels (drawn from a uniform distribution) to the original distance between the dot and line in one of the panels.

In same-different brightness detection blocks, the horizontal distance between the dot and the vertical line was always identical across the two panels within a trial (but varied across trials), and participants determined whether the brightness of the dot across the two panels was the same or different. In half of the trials in each block, the dots differed in brightness and participants indicated this by a button press. At the beginning of each block, a dummy trial with either a distance or brightness difference between the panels informed participants of the task in the upcoming block.

The same-different distance detection task identified cortical regions that process spatial relations between objects compared to regions that are sensitive to changes in brightness, which are anatomically distinct. The same-different brightness detection task, being visually similar to the distance task, (visually identical, apart from the within-trial differences in brightness) acted as a control to account for any activity that was not specifically related to distance processing per se, such as activity related to the shape and position of the stimuli in the display.

### Ventral stream localizer

In the ventral stream localizer tasks, we compared activations evoked by a same-different face detection task to activations evoked by a same-different house detection task, thereby identifying regions active more by faces compared to a non-face category of objects (the task was adapted from Kanwisher et al. 1997 and used previously in Zachariou et al. 2016). In separate blocks with counterbalanced order, the stimuli were either gray-scale images of faces or houses (22 images per category). In half of the trials of each block, the two items differed and participants indicated detection of this difference by button press (responses were withheld on matching trials). At the beginning of each block, a dummy trial consisting of two different faces or houses (presented for 1.7 s) informed participants of the task in the upcoming block.

### TBS sessions

Following the localizer session, participants completed three TBS sessions on different days. There were at least 24 hours between TBS sessions for safety reasons. In each TBS session, we delivered TBS on only one target site (left/right PPC and vertex; defined in the TBS protocol section of the Methods); the order of the sessions was counterbalanced across participants. Each TBS session comprised two fMRI scans, a pre-TBS scan and a post-TBS scan. Immediately after the pre-TBS scan, participants exited the MRI scanner and, within five minutes, TBS was delivered. Participants then returned to the MRI scanner and scanning resumed, on average, within three minutes and forty seconds after TBS to complete the post-TBS scan. Each TBS scan comprised four functional runs. A functional run consisted of twelve, 32-s blocks, each preceded and followed by 8 s of fixation. Six of these blocks comprised configural face difference detection trials (Figure 1A) and the remaining six comprised featural face difference detection trials (Figure 1B). These same/different face detection tasks are identical to the ones described in Zachariou et al. (2016).

**Figure 1.**
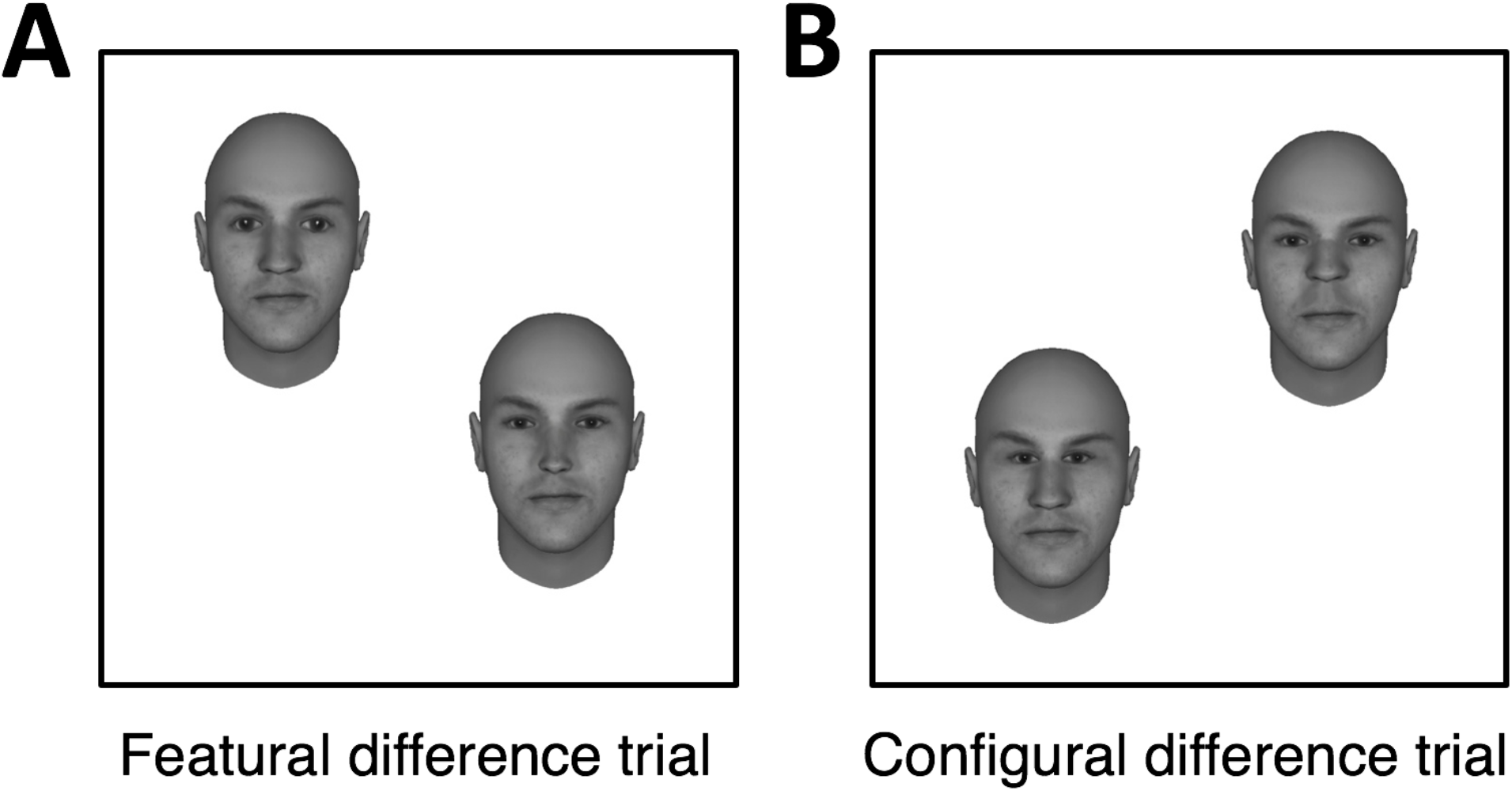
Sample trial displays and stimuli used during the TBS sessions. **A**. Sample trial display in which the exemplars differed in features (featural difference). **B**. Sample trial display in which the exemplars differed in configuration (configural difference).

Each trial lasted 2700 msec and trials within a block were separated by a 300-msec intertrial interval. On each trial, participants compared two faces, presented simultaneously on either side of the screen center, and indicated with a button press if they differed. If they did not differ, participants did not respond. Across trials, the two face images appeared in one of two possible spatial configurations: in half the trials, faces appeared in a top-left, bottom-right configuration, and in the remaining half, in a top-right, bottom-left configuration. The face stimuli were 4.5° wide by 5° tall and were separated by 4° of visual angle, 2° on either side of the screen center.

For the configural/featural face tasks of the TBS sessions, we used 20 different face exemplars, which were generated using the FaceGen software package. Following the creation of the face exemplars, we assigned two differently shaped sets of eyes and noses to each one of the 20 faces. Therefore, each face exemplar could appear in two variants, which differed only in the shape of the eyes and the nose. We will refer to the two variants of a face exemplar that differed in shape as S_1_ and S_2_.

Great care was taken in designing the stimuli of the featural task to ensure that the horizontal distance between the eyes (measured from the inside edge of both eyes) as well as the vertical distance between the top-most part of the mouth and the bottom-most part of the nose were identical between S_1_ and S_2_. We achieved this by manipulating the shape of the nose along the horizontal dimension and the shape of the eyes along the vertical dimension (maximum horizontal/vertical distance difference of 0.14 ° of visual angle between S_1_ and S_2_). This was done to minimize configural differences when shape was changed in the featural face task.

All face images were in gray-scale (each pixel only carried intensity information, a value from 0 – 100%). Using photoshop, we extracted the per-pixel intensity values of one face image (we used the “save image statistics” function in photoshop), which acted as the template image. The same procedure was used to extract the per-pixel intensity information from the areas of the template face image corresponding to the eyes and nose (separately for each face feature). We then applied these pixel intensity templates (using the same plugin) to every other face image and face feature used in the creation of the S_1_ and S_2_ variants. Consequently, every face image and face feature was very similar in luminance and contrast to the template face image. The average luminance value of the face images was 40.9%, the variance in luminance was 0.6% and the standard deviation was 0.75%.

In addition to the two sets of shape features, we also assigned two different spatial configurations to each face exemplar (the difference occurred in the distance between the eyes and between the nose and mouth; typical distance differences were between 0.5-0.9° of visual angle). Hence, the shape features (eyes and nose) of each face exemplar could appear with two different spatial configurations. We will refer to these two different spatial configurations as C_1_ and C_2_.

A featural difference occurred by presenting both the S_1_ and the S_2_ variants of a face exemplar on the stimulus display of a trial while holding spatial configuration (i.e. C_1_ or C_2_ randomly selected) constant between the face images. Similar to the featural-difference trials, configural-difference trials occurred when both the C_1_ and C_2_ spatial configurations of a face exemplar were presented within a stimulus display while holding S_1_ or S_2_ (randomly selected) constant between the face images.

In no-difference (identical face) trials, both faces on the stimulus display were assigned the same combination of features (S_1 or 2_) and configuration (C_1 or 2_), which were randomly selected. In each block, no difference was present in half the trials. Participants were never told about the two types of face differences and were only asked to make same-different judgments between faces.

### Training to match the featural/configural face tasks in difficulty

During the first session of the experiment (the training session) and at the beginning of each TBS session, participants were trained on the configural and featural face tasks using the following procedure: participants were first trained on the featural face task until their accuracy reached 90% correct. If the 90% accuracy threshold was not reached within 20 minutes, training stopped and the participant was excluded from the study. Participants were then trained on the configural face task and the difficulty level of the task was adjusted until 1) RT was within 100 msec of that on the featural face task and 2) the configural face task could be performed at the 90% accuracy threshold. Altering the distance between the eyes and between the nose and mouth controlled the difficulty level of the configural face task. Adjustments in difficulty were made after each training block of 10 trials and, if RT/accuracy did not match within 45 minutes, the participant was excluded from the study. We did not alter the difficulty level of the featural face task because, in order to manipulate difficulty, the shape of the face features would have needed to change, thereby altering the spacing between the face features and introducing a confound. In contrast, adjusting the difficulty level of the configural face task did not affect the shape features of a face.

### fMRI acquisition

Participants were scanned in a Siemens Skyra 3-Tesla scanner with a 32-channel head coil. Functional images were acquired with a multi-band (Feinberg and Setsompop, 2013), echo-planar imaging sequence (TR=2 s, TE=27 msec, flip angle 70°, 3.2 mm isotropic voxels, 72x72 matrix, field of view 205 mm, 72 axial slices covering the whole brain). The 72 slices were acquired with in-plane acceleration using the Siemens protocol iPAT with an acceleration factor of two. An MPRAGE sequence (1-mm^3^ voxels; 176 slices, field of view 256 mm) was used for anatomical imaging, and anatomical images were acquired in every scan session, including the localizer session.

The alignment of a participant’s brain between the pre- and post-TBS scans was accomplished using the Siemens auto-align function. This function first acquires a series of scout images (using 3D VIBE; a Siements T-1 weighted Gradient Echo Sequence), which are used to identify the anterior and posterior commissures (AC and PC), separately for each participant. Two auto-align scout scans were performed, one at the beginning of the pre-TBS scan and the other at the beginning of the post-TBS scan. Since the anatomy of a participant’s brain did not change between the two TBS scans, the AC and PC identified in each scan had the same position relative to the whole brain, irrespective of the overall position of the head. The 72 EPI slices of each run of a TBS scan were then centered on the AC/PC plane identified by the auto-align scouts, thereby ensuring that all EPI slices, irrespective of scan (pre- or post-TBS), shared the same location relative to a participant’s AC/PC plane.

### fMRI Pre-processing

The functional scans were slice scan-time corrected, motion corrected, co-registered to their constituent contrast-corrected anatomical images (both the functional and anatomical scans were corrected for contrast artifacts during acquisition using the Siemens pre-scan normalize function), normalized to Talairach space (Talairach and Tournoux 1988) using a non-linear transformation (3dQwarp; http://afni.nimh.nih.gov/pub/dist/doc/program_help/3dQwarp.html), smoothed with a Gaussian kernel of 6 mm FWHM, and mean-based intensity normalized (all volumes by the same factor) using AFNI (Cox 1996). In addition, linear and non-linear trends were removed during pre-processing of the data.

Additional pre-processing steps were performed prior to the functional connectivity analysis, according to the basic ANATICOR regression-based approach (e.g. Jo et al. 2010; Gotts et al. 2012; Stoddard et al. 2016). Using each participant’s anatomical scan (MPRAGE), ventricles, gray and white matter segmentation masks were created (using SPM12; http://www.fil.ion.ucl.ac.uk/spm/software/spm12/), separately for each participant. All masks were resampled to the EPI voxel resolution, and ventricle and white matter masks were eroded by one voxel (or two voxels if task-like components were observed in the first three principal components of the PCA analysis discussed below) to prevent partial volume effects with gray matter. We then extracted separate nuisance time series for ventricles and white matter. In total, the nuisance regression for each participant involved 11 regressors of no interest: six motion parameters, one average ventricle time series, one localized estimate of white matter (averaging within a sphere of radius 20 mm centered on each voxel), and the first three principal components of all voxel time series from a combined ventricle and white matter mask, calculated after first detrending with AFNI’s fourth-order polynomial baseline model (as in Stoddard et al. 2016; comparable to aCompCor: Behzadi et al. 2007). After this nuisance model was subtracted from each participant’s EPI data to obtain the cleaned residual time series, a task regression was performed to further remove any evoked responses from the blocks during the task (using the BLOCK model in AFNI’s 3dDeconvolve). The resulting time series were then extracted separately from blocks of different conditions, with blocks of the same type concatenated together for purposes of condition comparisons after adjusting for the delay in the BOLD signal in each block (6 s after the start of each block until 4 s after the end). Estimates of the level of residual global artifacts present in the residual time series (including factors such as head motion, cardiac and respiration effects, etc.) were calculated per condition for later use as nuisance covariates in group-level analyses using the global level of correlation or “GCOR” (e.g. Gotts et al. 2013; Saad et al. 2013), which is the grand average correlation of all gray matter voxels with each other.

### fMRI statistical analyses

All imaging data were analyzed using AFNI (Cox 1996). Data from the localizer and TBS sessions were analyzed using linear mixed-effects models (*3dLME*; Chen et al. 2013). The resulting statistical maps were thresholded at qFDR < 0.05 using the false discovery rate approach for multiple comparisons correction (Genovese, Lazar, and Nichols, 2002). The motion parameters from the output of the volume registration step were regressed out in all AFNI analyses.

Seed-to-whole-brain functional connectivity analyses using the right aIT, FFA and OFA as seeds were performed as follows. First, brain masks were created for each participant that comprised gray matter voxels only (using SPM12). Then, we expanded spheres with a radius of 7 mm (60 voxels per sphere) positioned at the center of mass of each individually-defined right aIT, right FFA and right OFA. These 60 voxel-wide spheres acted as the seed ROIs for the functional connectivity analysis. Time series were extracted from these seed ROIs and then correlated (Pearson’s r) with the cleaned residual time series in all gray matter voxels (described in the pre-processing section). Correlation maps for each seed ROI were calculated separately for featural and configural face difference detections, TBS session (left/right PPC, vertex), TBS scan (pre-, post-TBS) and participant. The maps were then transformed using Fisher’s z to yield normally distributed values. A group-level analysis was then performed, using linear mixed-effects models (*3dLME*; Chen et al. 2013), using the correlation maps from each of the above conditions as factors, both in the entire gray matter mask as well as in a mask constructed by the localizer-defined ROIs (bilateral PPC, aIT, FFA, OFA and PPA). In order to rule out contributions of global BOLD artifacts (e.g. head motion, cardiac/respiration fluctuations) to the condition differences, GCOR (described in the pre-processing section) was added as a nuisance covariate to these condition comparisons (Gotts et al. 2013).

Behavioral data collected during the scans were analyzed using SPSS and a general linear mixed-effects model (with participants added as a random variable). Multiple comparisons used Sidak corrections where necessary.

### Eye tracking

Eye tracking was conducted outside the MRI scanner using an SR Research (Ottawa, ON, Canada) EyeLink 1000 Plus eye-tracking system and a desktop monitor (BENQ XL) with the same resolution, physical dimensions of the viewable area of the screen and distance from the participants’ eyes as the projection screen inside the MRI scanner (450 × 344 mm viewable area on the screen, 29° visual angle horizontally by 18° vertically at a distance of 128 cm away from the participants’ eyes). Eye-tracking data were collected during the training session, immediately after successful completion of training. During this eye-tracking section, participants completed two functional runs, identical to those of the TBS sessions. The SR Research data viewer software was used to define areas-of-interest (AOIs) and to extract number of fixations and fixation durations (in msec) from each AOI. We then imported these eye-fixation data to SPSS for statistical analyses. The following procedure was used to compare the pattern of eye fixations between the configural and featural difference detection tasks across trials. For each of the four on-screen locations where face images appeared (two locations per trial), 16 AOIs were constructed in a 4x4 matrix, covering the entire region occupied by a face image (64 AOIs in total). The width of all AOIs was 1.0° of visual angle and their height was 1.5°.

The number of eye fixations and fixation durations in each AOI, for each type of face difference, were calculated for each participant. ANOVAs were then conducted separately for each dependent measure with type of task (featural/configural face difference detections) and AOI (1-64) as factors.

### TBS Protocol

TBS was delivered using a continuous TBS paradigm (Huang et al., 2005) as a train of three biphasic (equal amplitude) TMS pulses at 50 Hz, repeated at 5 Hz, for a total of 900 pulses or 300 thetabursts. TBS was delivered at 30% of the maximum stimulator output (on average at about 80% of active motor threshold; see Pitcher et al. 2017). Pulses were delivered using a figure-eight coil (50 mm external diameter) in conjunction with a Magstim Super Rapid 2 stimulator (Magstim; Whitland, UK). Target sites for TBS were selected bilaterally and corresponded to the peak activity voxels within left and right PPC identified using the visuospatial localizer task (from the dorsal stream localizer) and the fMRI contrast of activations evoked by same/different distance detections > activations evoked by same/different brightness detections. These target sites were selected at the individual subject level. The vertex, the topmost center part of a participant’s head, defined as the midpoint between the inion and the nasion of the head, was the control site. TBS sites were targeted in real time using the Brainsight system (Rogue Research).

## Results

### Localizing ROIs

We first localized brain regions within the dorsal visual pathway that mediate spatial perception and regions within the ventral visual pathway that mediate face perception. These regions-of-interest (ROIs) were identified as follows: 1) activations during a same/different distance detection task > those during a same/different brightness detection task identified visuospatial processing regions in PPC; and 2) activations during a same/different face detection task > those during a same/different house detection task identified face processing regions within the ventral visual pathway. All ROIs were identified at both the group and the individual participant level at a significance threshold of qFDR < 0.05.

For the PPC, all positively activated regions within each hemisphere were used to create masks that served as the dorsal stream/PPC ROIs. At the group level, positive activations included bilateral precuneus (BA7; center of mass Talairach coordinates on the right: 16, −67, 49; center of mass Talairach coordinates on the left: −17, −66, 49), right superior parietal lobule (center of mass Talairach coordinates: 25, −58, 57) and right inferior parietal lobule (BA 40; center of mass Talairach coordinates: 35, −39, 42).

For the VOTC, at the group level, positively activated regions included bilateral OFA in the middle occipital gyrus (within 1 mm of BA 19; center of mass Talairach coordinates on the right: 43, −71, −7; center of mass Talairach coordinates on the left: −38, −78, −10), bilateral FFA in mid-fusiform gyrus (BA 37; center of mass Talairach coordinates on the right: 41, −43, −19; center of mass Talairach coordinates on the left: −38, −47, −19), and bilateral aIT in anterior inferior temporal gyrus (BA 21; center of mass Talairach coordinates on the right: 32, −6, −31; center of mass Talairach coordinates on the left: −28, −9, −26).

In addition to these face-selective ROIs, we defined two bilateral ROIs within the parahippocampal gyrus, corresponding to the parahippocampal place area (PPA; Epstein and Kanwisher 1998). These PPA ROIs showed greater activations evoked by same/different house detections than same/different face detections (BA36; center of mass Talairach coordinates on the right: 27, −40, −9; center of mass Talairach coordinates on the left: −25, −43, −9). The PPA ROIs were used to explore whether the effects of TBS were specific to face-selective ROIs in VOTC or also included scene-selective regions in the parahippocampal cortex.

### TBS sessions

Three TBS scan sessions followed the localizer scan session. In each, participants performed same/different featural and configural face detection tasks (see Methods and Figures 1A and 1B).

To evaluate the effects of TBS, we subtracted the fMRI activity recorded post-TBS from that recorded pre-TBS, separately for the different tasks (configural/featural face difference detections) and different TBS sites (right/left PPC and vertex). Activity from each pre-TBS scan served as a baseline for the post-TBS scans, revealing the effect of TBS. Since the tasks performed pre- and post-TBS were identical, each face task (configural/featural) acted as its own control.

### Effect of TBS on magnitude of activation

We first investigated the effect of TBS on the magnitude of activity (% signal change) at the level of the whole brain, but this analysis did not yield any significant main effects or interactions between the factors. We therefore repeated the analysis within individually-defined VOTC ROIs. ANOVAs were conducted using a general linear mixed effects model with participants added as a random variable. The pre-post TBS percent signal change in activity (extracted from the individual subject ROIs) was the dependent measure and task type (configural/featural difference detections), TBS target site (right PPC, left PPC and vertex) and ROI (left/right OFA, FFA, aIT and PPA) were the factors. None of the main effects of this analysis were significant (ROI: F(7,16) = 1.47, p = 0.17; TBS site: F(2,16) = 1.67, p = 0.19; task type: F(1,16) = 1.9, p = 0.1). Similarly, the two-way interactions between ROI and TBS site (F(14,16) = 0.82, p = 0.65) and between ROI and task type (F(7,16) = 0.75, p = 0.63) were also not significant. There was, however, a significant two-way interaction between task type and TBS site (F(2,16) = 4.8, p = 0.01) and a significant three-way interaction between task type, TBS site and ROI (F(14,16) = 4.6, p = 0.04). To unpack the significant three-way interaction between task type, TBS site and ROI, we conducted separate analyses for each TBS target site with task type and ROI as factors.

The analysis in the left PPC target site yielded a significant interaction between task type and ROI (F (7, 16) = 2.70, p = 0.01; Figure 2A). To evaluate this significant interaction, we conducted separate analyses for each ROI in VOTC, with task type as the sole factor. The results indicated that in right OFA (F(1,16) = 7.3 p = 0.01) and right aIT (F(1,16) = 16.1, p = 0.009) task type was a significant main effect; in both areas, the pre-post difference in fMRI activity was greater for configural than for featural face difference detections (right OFA: face configural = 0.17%, face featural = −0.09%; right aIT: face configural = 0.11%, face featural = −0.03%). The analyses on the remaining ROIs did not yield task type as a significant main effect (right FFA: F(1, 16) = 2.08, p = 1.7; right PPA: F(1,16) = 0.64, p = 0.44; left OFA: (F1, 16) = 0.003, p = 0.96; left FFA: F(1, 16) = 0.19, p = 0.67; left aIT: F(1, 16) = 0.001, p = 0.99; left PPA: F(1, 16) = 0.22, p = 0.64). Thus, TBS delivered over the left PPC caused a significantly larger decrease in activity for configural than featural face difference detections in right OFA and right aIT only.

**Figure 2.**
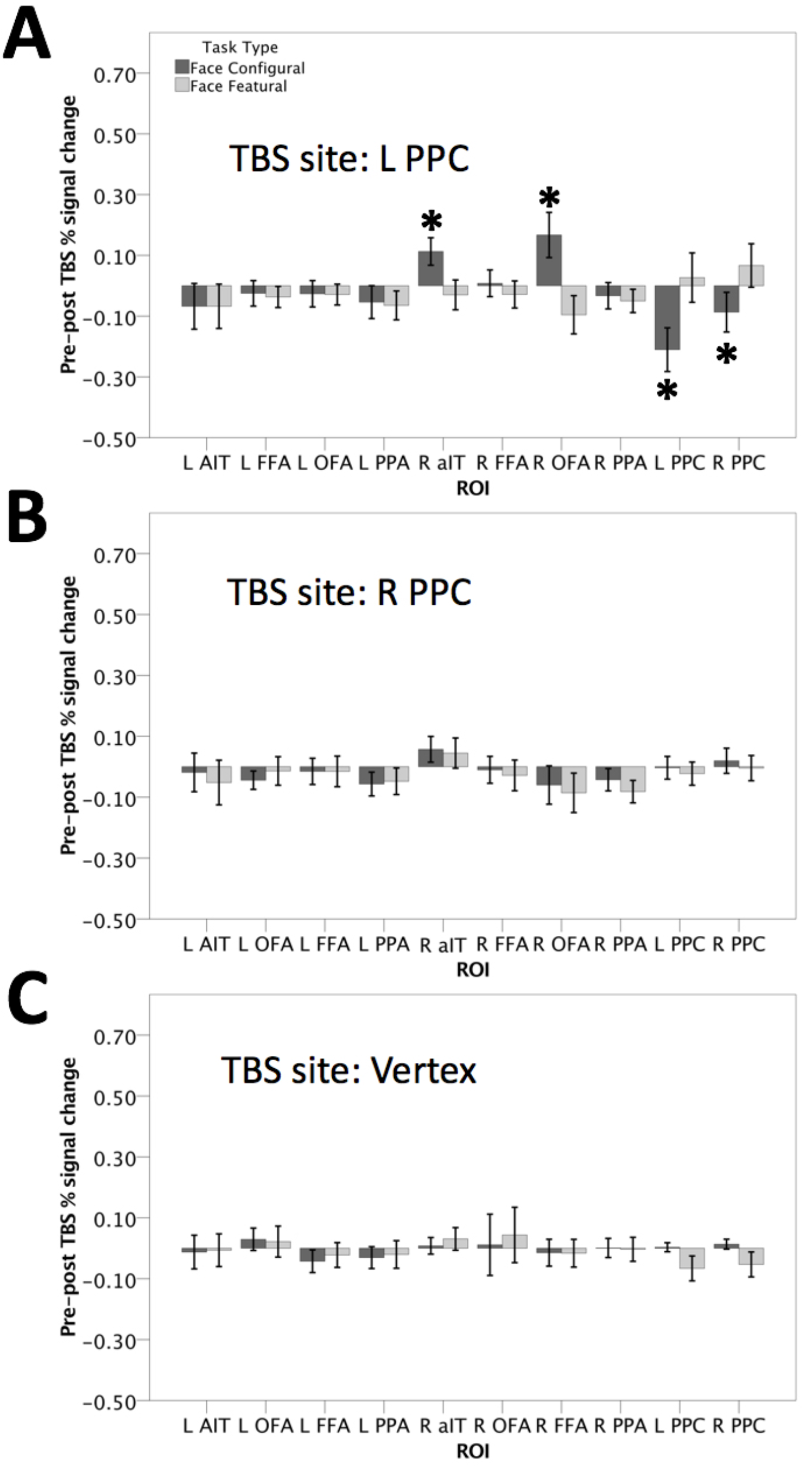
The panels depict the pre-minus post-TBS percent signal change in fMRI activity for configural and featural face difference detections within all ROIs used in the experiment: bilateral PPC, aIT, FFA and OFA, and for all TBS target sites: right PPC, left PPC and vertex. **A**. Pre-minus post-TBS percent signal change in fMRI activity for configural and featural face difference detections, across ROIs, when TBS was delivered on left PPC. The asterisk (*) indicates those ROIs where the pre-post activity for configural face difference detections was significantly different from that of featural face difference detections. **B**. Pre-minus post-TBS percent signal change in fMRI activity for configural and featural face difference detections, across ROIs, when TBS was delivered on right PPC. **C**. Pre-minus post-TBS percent signal change in fMRI activity for configural and featural face difference detections, across ROIs, when TBS was delivered on the vertex control site. The error bars denote +/-1 SEM, n = 17.

To investigate whether these differences we observed in the magnitude of activity between the pre- and post-scans for TBS on the left PPC were significantly different from zero, we conducted one-sample t-tests (two-tailed; against zero) on the pre-post TBS percent signal change, separately for configural and featural face difference detections, within each ROI. For configural face difference detections, the pre-post TBS activity was significantly different from zero in right OFA (t(16) = 2.25, p = 0.04) and right aIT (t(16) =2.78, p = 0.01). In the remaining face-selective ROIs, the pre-post TBS activity was not significantly different from zero: right FFA (t(16) =0.18, p = 0.86); left aIT (t(16) =-0.90, p = 0.38); left FFA (t(16) = −0.6, p = 0.56); left OFA (t(16) =-0.61, p = 0.55). Similarly, the pre-post TBS activity within bilateral PPA was not significantly different from zero: right PPA (t(16) =-0.76, p = 0.46); left PPA (t(16) = −0.99, p = 0.33).

For featural face difference detections, the pre-post TBS activity for the left PPC target site was not significantly different from zero in any of the VOTC ROIs: right aIT (t(16) = −0.77, p = 0.45); right FFA (t(16) = −0.65, p = 0.52); right OFA (t(16) = −1.52, p = 0.15); right PPA (t(16) = −1.30, p = 0.21); left aIT (t(16) = −0.93, p = 0.37); left FFA (t(16) = −1.06, p = 0.31); left OFA (t(16) = −0.84, p = 0.41); left PPA (t(16) = −1.36, p = 0.19). Thus, TBS delivered on the left PPC decreased activity for configural but not for featural face difference detections. This effect was confined to the right OFA and right aIT. In all other VOTC ROIs, including bilateral PPA, activity did not differ between the pre- and post-TBS scans, irrespective of task type.

The analysis for the right PPC (Figure 2B) and vertex (Figure 2C) TBS sites did not yield any significant main effects or interactions between the factors: right PPC: task type: F(1, 16) = 0.42, p = 0.52; ROI: F(7,16) = 0.40, p = 0.5; task type x ROI: F(7,16) = 0.22. p = 0.98; vertex: task type: F(1, 16) = 0.28, p = 0.93; ROI: F(7,16) = 0.60, p = 0.75; task type x ROI: F(7,16) = 0.07. p = 0.99.

Following the analyses on task type (configural/featural face difference detections), which were performed separately for each TBS site and ROI, we conducted additional ANOVAs, in which separate analyses were conducted for each level of task type (featural and configural face processing) with TBS target site (left/right PPC and vertex) and ROI (left/right OFA, FFA, aIT and PPA) as the main factors. These analyses evaluated whether the pre-post TBS activity for configural and featural face difference detections differed between the PPC TBS target sites and the vertex control site.

For configural face difference detections, we observed a significant interaction between ROI and TBS site (F(14,16) = 2.6, p = 0.02). To evaluate this significant interaction, we repeated the analysis separately for each ROI with TBS site as the sole factor. We found that TBS site was a significant main effect in only two of the VOTC ROIs: right aIT (F(2,16) = 3.95, p = 0.03) and right OFA (F(2,16) = 3.91, p = 0.03). For right aIT, pair-wise comparisons (adjusted for multiple comparisons using Sidak) between the three TBS sites indicated that, for the left PPC target site, pre-post TBS activity differed significantly from both the vertex control site (p = 0.03) and the right PPC target site (p = 0.04). For the right PPC target site, pre-post TBS activity did not differ significantly from the vertex (p = 0.48). For the right OFA ROI, pair-wise comparisons indicated that the pre-post activity for the left PPC target site also differed significantly from both the vertex site (p = 0.04) and the right PPC target site (p = 0.03). The comparison of the pre-post TBS activity between the right PPC target site and the vertex site was not significant (p = 0.78). For the remaining ROIs, TBS site was not a significant main effect: right FFA (F(2,16) = 1.05, p = 0.90), right PPA (F(2,16) = 0.37, p = 0.69), left aIT (F(2,16) = 0.31, p = 0.74), left FFA (F(2,16) = 0.13, p = 0.88), left OFA (F(2,16) = 1.07, p = 0.35), left PPA (F(2,16) = 0.12, p = 0.88).

For featural face difference detections, there were no significant main effects or a significant interaction between the factors (TBS site: F(2,16) = 2.16, p = 0.08; ROI: F(7,16) = 0.431, p = 0.88; TBS site x ROI: F(14,16) = 0.41, p = 0.97).

In summary, the results indicated that the effect of TBS on the magnitude of fMRI activity was significantly stronger when delivered on the left PPC target site than the vertex control site or the right PPC site, but only for configural face difference detections and only in right OFA and right aIT.

We next repeated the same analyses on the PPC ROIs in order to evaluate the effects of TBS on the target sites. The ANOVA with task type (configural/featural face difference detections), TBS site (left/right PPC and vertex) and ROI (left/right PPC) as factors yielded a significant two-way interaction between task type and TBS site (F(2,16) = 10.1, p < 0.001). The three-way interaction between task type, TBS site and ROI was not significant (F(2,16) = 0.29, p = 0.75). To unpack the significant two-way interaction between task type and TBS site, we repeated the analysis separately for each TBS site.

The analysis of the left PPC TBS site yielded task type as the only significant main effect (F(1,16) = 9.92. p < 0.001). TBS on the left PPC caused a significantly larger increase in the magnitude of activity for configural (-0.15%) than featural face difference detections (0.04%); this effect was not statistically different between the left and right PPC ROIs. Thus, like the results for the VOTC ROIs, TBS on left PPC affected fMRI activity more during configural than featural face difference detections.

The analysis of the right PPC and vertex TBS sites yielded no significant main effects or interactions between the factors: right PPC: task type: F(1, 16) = 1.9, p = 0.18; ROI: F(1,16) = 1.7, p = 0.2; task type x ROI: F(1,16) = 0.02. p = 0.88; vertex: task type: F(1, 16) = 1.3, p = 0.83; ROI: F(1,16) = 0.17, p = 0.68; task type x ROI: F(1,16) = 0.003. p = 0.96. Thus, like the findings in right VOTC, there was a differential effect in left PPC between changes in activity for configural and featural face difference detections, but only when TBS was delivered on the left PPC. Also in accord with the findings in VOTC, the magnitude of the pre-post TBS effect was significantly greater for configural compared to featural face difference detections. However, when TBS was delivered on the left PPC, fMRI activity during configural face difference detections decreased in the face-selective ROIs of right aIT and right OFA, whereas activity for the same task increased in the spatial processing ROIs of PPC.

We then performed separate analyses for each level of task type (featural and configural face processing) with TBS target site (left/right PPC and vertex) and ROI (left/right PPC) as the main factors. These analyses evaluated whether the pre-post TBS activity for configural and featural face difference detections differed between the TBS PPC sites and the vertex control site. For configural face difference detections, we observed a significant main effect of TBS site only (F(2,16) = 9.53, p < 0.001). Pair-wise comparisons between the TBS sites indicated that, for the left PPC target site, the pre-post TBS activity was significantly greater (-0.15) than both the right PPC target site (0.08; p < 0.001) and the vertex site (0.08; p < 0.001). In contrast, TBS on the right PPC target site did not differ significantly from TBS on the vertex site (p = 1.00). The main effect of ROI was not significant (F(1, 16) = 2.37, p = 0.13) and there was no significant interaction between ROI and TBS site (F(2, 16) = 1.12, p = 0.33).

For featural face difference detections, there were no significant main effects or a significant interaction between the factors (ROI: F(1, 16) = 0.36, p = 0.55; TBS site: F(2, 16) = 2.4, p = 0.1; ROI x TBS site: F(2, 16) = 0.04, p = 0.96). Thus, in accord with all previous findings, only TBS on the left hemisphere differed from TBS on the vertex control site and only during configural face difference detections.

### Why was TBS on right PPC ineffective?

The absence of a TBS effect from the right PPC site was puzzling and could indicate either that: 1) the processing of spatial information related to configural face processing is lateralized to the left hemisphere; or 2) TBS on the right hemisphere failed to produce measurable differences in fMRI activity. The first possibility is inconsistent with our previous findings of bilateral PPC activation for configural compared to featural face processing and impaired behavioral performance on configural face difference detections when rTMS was delivered on either the left or right PPC (Zachariou et al. 2016). We therefore believe the second possibility is more likely; namely, that TBS may not have been sufficiently effective over the right PPC to modulate its activity. We believe this second possibility is more likely for two reasons: 1) based on previous research (Nummenmaa et al. 2013; Silson et al. 2013; Strong et al. 2017) the area of stimulation using a 50 mm, figure-eight TMS coil at 60-65% machine output is about 10 mm^2^. The area of stimulation is presumably smaller at 30% machine output, which is approved for use with TBS by the NIMH IRB. 2) According to our previous study (Zachariou et al. 2016 and also replicated below) the activation in response to configural face difference detections covers a disproportionally larger area of cortex in the right compared to the left hemisphere PPC. Consequently, it is possible that the area of cortex stimulated by TBS may not have been sufficiently large to interfere with the magnitude of configural face processing activity in the right PPC but sufficiently large to interfere with the magnitude of configural face processing activity in the left PPC due it’s smaller area.

To explore this possibility, we examined the activity evoked by configural and featural face difference detections in a whole-brain, group-level analysis (at qFDR < 0.05). For this analysis, we focused on activity from the pre-TBS scans only to exclude TBS-induced differences in the magnitude of activation for the two face tasks. As such, TBS session (left/right PPC and vertex) and task type (configural/featural face difference detections) were the factors. The analysis yielded a main effect of task type. As anticipated, TBS session was not a significant main effect and there was no interaction between the factors (up until qFDR = 0.95), indicating that the magnitude of activity for configural and featural face difference detections did not change significantly between TBS sessions (scanning participants on different days) for the pre-TBS scans. To further explore the significant main effect of task type, we used the fMRI contrast of activations evoked by configural difference detections > those evoked by featural difference detections, also thresholded at qFDR < 0.05 (Figure 3).

**Figure 3.**
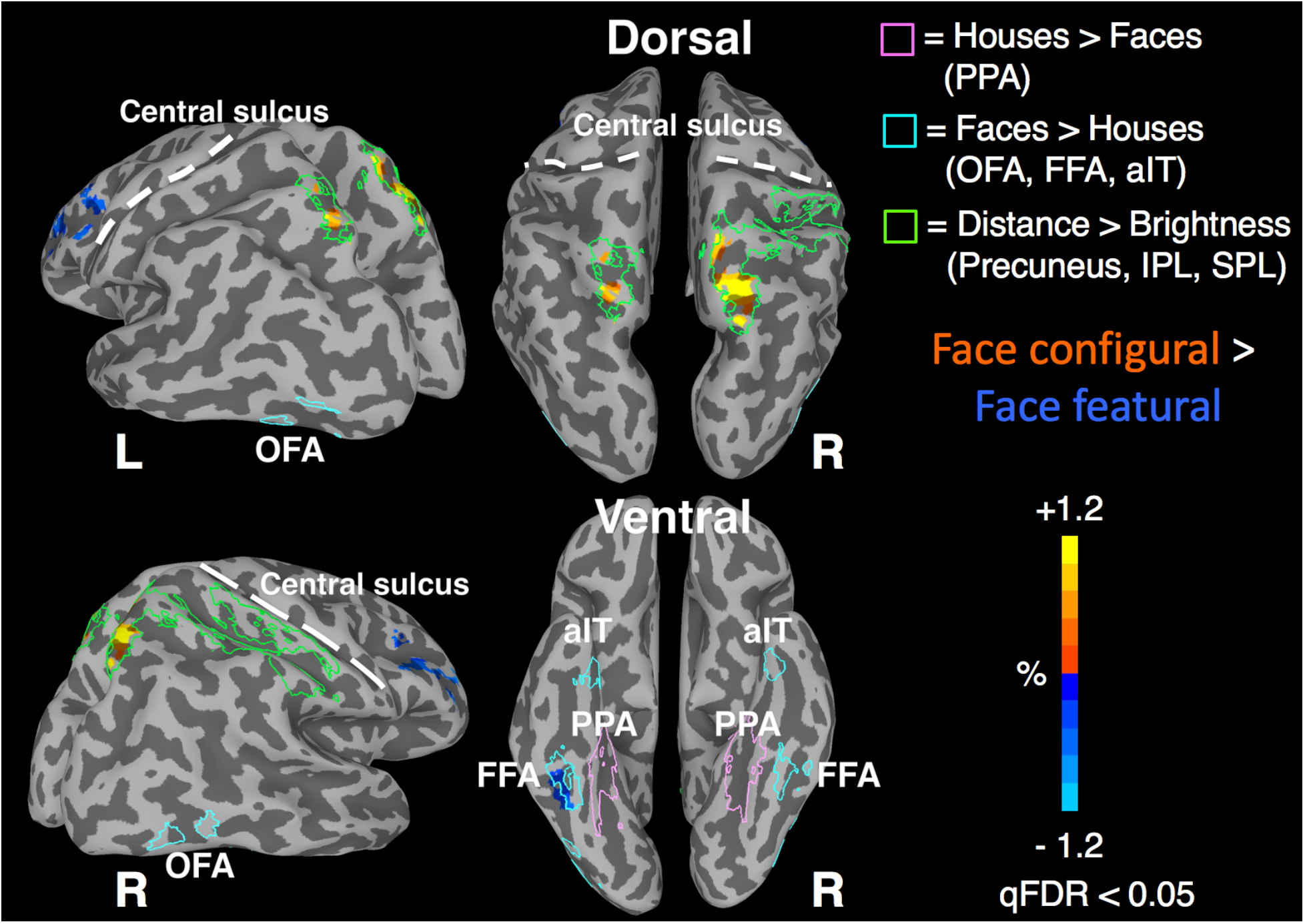
Cortical activations for configural and featural face difference detections at the level of the whole brain. The figure depicts brain activity from the fMRI contrast of configural face difference detections > featural face difference detections, calculated from a group level, whole brain analysis of the pre-TBS scans merged across TBS session. Positive activations (orange to yellow) correspond to regions more active for configural face difference detections, whereas negative activations (cyan-blue) correspond to regions more active for featural face difference detections. The green outlines illustrate the brain regions identified by the visuospatial localizer task (distance difference detections > brightness difference detections). The regions identified by this localizer were the bilateral precuneus, the right inferior and superior parietal lobules (IPL, SPL). The cyan outlines illustrate the brain regions identified by the face localizer task (face difference detections > house difference detections). The regions identified by this localizer were bilateral aIT, FFA and OFA. The pink outlines illustrate regions more active in response to house difference detections compared to face difference detections and consist of bilateral PPA. For clarity, the anatomical locations of the central sulci are shown with dashed white lines. n = 17.

This fMRI contrast revealed two positively activated clusters in PPC, which overlapped considerably with the localized spatial processing ROIs used to define the TBS targets. The cluster of activity within the left PPC (31 voxels) was within the precuneus (center of mass Talairach coordinates: −18, −68, 49; peak voxel Talairach coordinates: −19, −70, 50), whereas the cluster within the right PPC was more than three times larger (116 voxels), covering regions in the precuneus and superior parietal lobule (peak voxel Talairach coordinates: 14, −73, 53 within the right precuneus; center of mass Talairach coordinates: 14, −68, 53 in the right superior parietal lobule). These findings support the possibility that TBS with a stimulation area of < 10mm^2^ affected a much larger portion of the configural face activity in the left PPC than the right PPC.

### Effect of TBS on task-based functional connectivity

We next explored whether TBS also affected the level of task-based functional connectivity between the localized ROIs in VOTC and PPC. We used three seed ROIs: a 7-mm radius sphere (60 voxels per sphere) positioned at the center of mass of the right aIT, right FFA and right OFA ROIs defined at the individual participant level (see Methods for details). For each seed, we generated functional connectivity maps, at the individual participant level, between that seed and the rest of the brain, separately for each task type (configural/featural face difference detections), target site (left/right PPC, vertex) and TBS scan (pre-/post-TBS). The resulting functional connectivity maps were used as input to a linear mixed effects model (*3dLME*; Chen et al. 2013) with task type, target site, TBS scan and seed as factors. All resulting statistical maps were adjusted for multiple comparisons using the false discovery rate at qFDR < 0.05.

At the level of the whole brain, the results from the functional connectivity analysis did not survive the multiple comparisons correction. We therefore repeated the analysis within a smaller mask of the previously localized ROIs (namely, bilateral aIT, FFA, OFA, PPA and PPC, defined at the group level) in order to reduce the number of statistical comparisons to those voxels of most interest. This analysis yielded a significant four-way interaction between task type, target site, TBS scan and seed. To unpack this interaction, we repeated the analysis separately for each target site.

For both the left and right PPC sites, there was a significant three-way interaction between task type, TBS scan and seed. In contrast, for the vertex control site, there were no significant main effects or interactions between the factors at the same significance threshold. To explore whether TBS on the left and right TBS sites differed, we repeated the initial analysis but excluded the vertex TBS site from the factors. This analysis yielded a significant three-way interaction between task type, TBS scan and seed but the four-way interaction between the factors was not significant. Consequently, unlike the analyses on the magnitude data, the effect of TBS on the degree of functional connectivity was comparable between the left and right PPC targets. To further explore the significant three-way interaction between task type, TBS scan and seed, we repeated the analysis separately for each seed ROI.

For the right aIT seed, we observed a significant two-way interaction between task type and TBS scan. To explore this two-way interaction, we used the following functional contrasts: activations to face configural pre-TBS > those to face configural post-TBS and activations to face featural pre-TBS > those to face featural post-TBS (Figure 4). The configural pre > post-TBS contrast yielded positive activations within bilateral FFA, bilateral OFA and bilateral PPC ROIs. Thus, for configural face difference detections, the level of functional connectivity between the right aIT and bilateral FFA, bilateral OFA and bilateral PPC ROIs decreased following TBS over either the left or right PPC targets; the degree of functional connectivity between the right and left aIT and between the right aIT and bilateral PPA was not affected by TBS.

**Figure 4.**
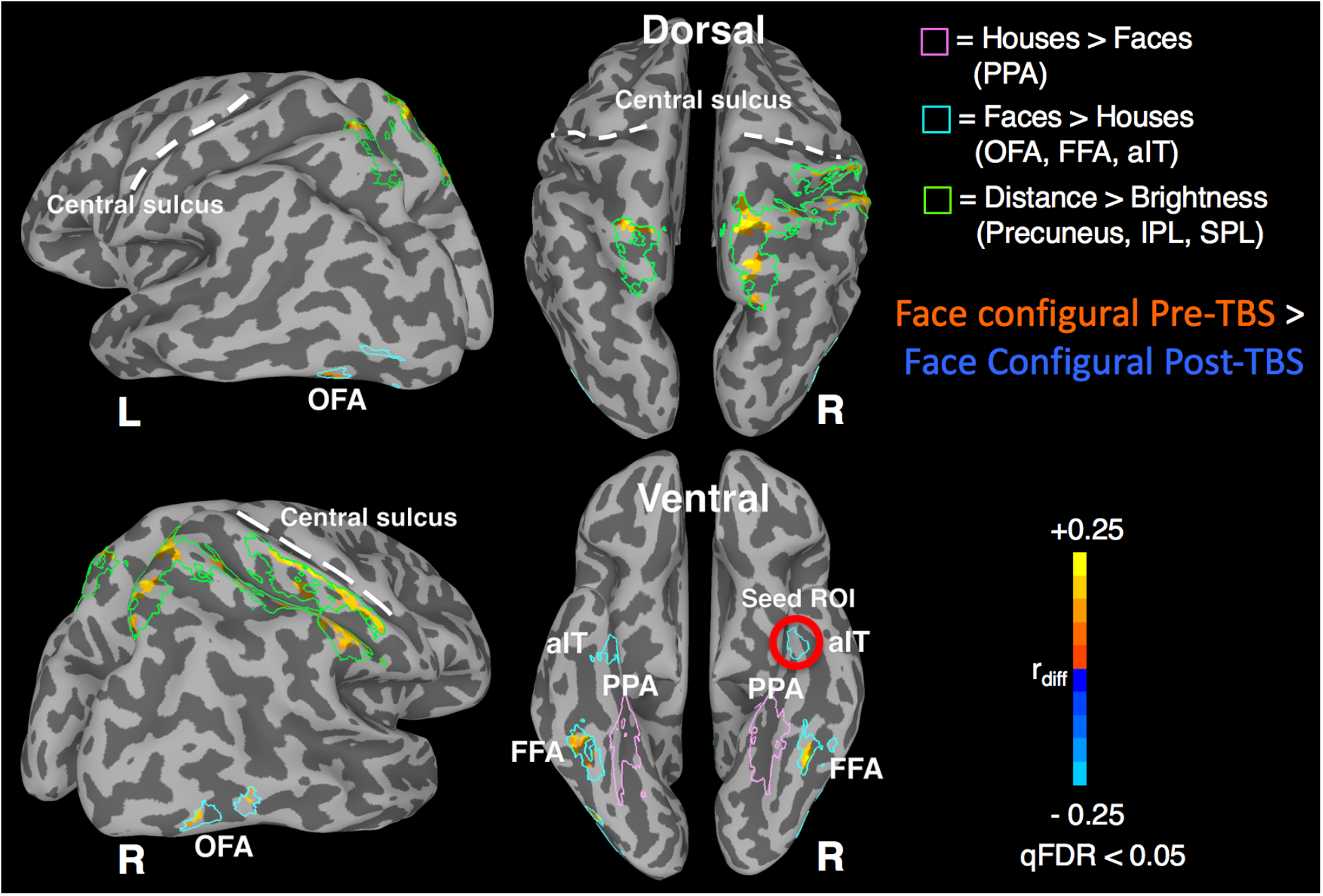
Differences in functional connectivity (differences in Pearson’s r) between the pre- and post-TBS scans for configural face difference detections, when TBS was delivered on either the left or right PPC. Differences in the level of functional connectivity were explored between the right aIT as the seed and the following ROIs: left aIT, bilateral FFA, bilateral OFA, bilateral PPC and bilateral PPA (bilateral PPA acted as control ROIs). All brain regions in which the level of functional connectivity between them and the seed ROI decreased post-TBS compared to pre-TBS (for configural face difference detections only) are depicted as positive activations (orange-yellow). Brain areas where the reverse was true would be depicted as negative activations (cyan-blue). The green outlines illustrate the brain regions identified by the visuospatial localizer task (distance difference detections > brightness difference detections). The regions identified by this localizer were bilateral precuneus, right IPL and SPL. The cyan outlines illustrate the brain regions identified by the face localizer task (face difference detections > house difference detections). The regions identified by this localizer were bilateral aIT, FFA and OFA. The pink outlines illustrate regions more active in response to house difference detections compared to face difference detections and consist of bilateral PPA. For clarity, the anatomical locations of the central sulci are shown with dashed white lines. n = 17.

For featural face difference detections, there were no significant changes in functional connectivity in any of the VOTC ROIs. Consequently, TBS on the left and right PPC targets sites affected the degree of functional connectivity for configural face difference detections but not for featural face difference detections.

The functional connectivity analyses on the right FFA and right OFA seed ROIs did not yield any significant main effects or interactions between the factors, so we did not explore these further.

### Behavioral performance

To ensure that behavioral performance for featural and configural face difference detections was comparable, we trained participants on the face tasks first during the initial training session and then again prior to each TBS session so that RT and accuracy on the two tasks were matched across participants (see Methods). To evaluate whether the matching on task difficulty was successful while participants performed the tasks inside the MRI scanner, we conducted a repeated measures ANOVA with task type (configural/featural difference detections) and TBS target (left/right PPC and vertex) as factors, focusing only on behavioral data from the pre-TBS scans. Focusing on the pre-TBS scans only, excluded the effects of TBS on behavioral performance allowing us to evaluate how well the two tasks were matched following training. Separate analyses were conducted for RT and accuracy.

The analysis of the accuracy data yielded no significant main effects (TBS site: F(2,16) = 2.0, p = 0.14; task type: F(1,16) = 0.51, p = 0.60) or a significant interaction between the factors (TBS site X task type F(2,16) = 0.07, p = 0.94). For the pre-TBS scan sessions, the accuracy of the configural face task (89% accurate) was comparable to that of the featural face task (88% accurate) irrespective of TBS site.

The analysis of the RT data yielded similar results to those of the analysis on accuracy: no significant main effects (TBS site: F(2,16) = 0.7, p = 0.36; task type: F(1,16) = 0.8, p = 0.38) or a significant interaction between the factors (TBS site X task type F(2,16) = 0.39, p = 0.68). During the pre-TBS scans, participants were comparably fast in responding to configural (1429 msec) and to featural (1436 msec) face difference detections, irrespective of TBS site.

As expected, training the participants and adjusting the difficulty level of the configural face task, successfully matched the participants’ behavioral performance on the two face tasks. Following the analyses of the behavioral data from the pre-TBS scans only, we expanded the ANOVA to include behavioral data from both the pre- and post-TBS scans by adding TBS scan (pre- or post-TBS) as a factor. This analysis was conducted in order to evaluate the effects of TBS on behavioral performance. Separate analyses were conducted for RT and accuracy.

The analysis of the accuracy data yielded TBS scan as the only significant main effect (F(1,16) = 97.5, p < 0.001). Irrespective of TBS target site or task type, participants were slightly less accurate during the pre-TBS scan (89% correct) than the post-TBS scan (92% correct). All remaining main effects and interactions between the factors were not significant: TBS site: F(2,16) = 1.53, p = 0.22; task type: F(1,16) = 0.81, p = 0.38; TBS site X TBS scan: F(2,16) = 0.29, p = 0.74; TBS site X task type: F(2,16) = 0.91, p = 0.40; TBS session X task type: F(1,16) = 0.08, p = 0.78; TBS site X TBS scan X task type: F(2,16) = 1.16, p = 0.31).

The analysis of RT also yielded TBS scan as the only significant main effect (F(1,16) = 54.1, p < 0.001). Participants were slightly faster to respond during the post-TBS scan (1391 msec) than the pre-TBS scan (1432 msec), irrespective of TBS target site or task type. Like the accuracy results, all remaining main effects and interactions between the factors were not significant: TBS site: F(2,16) = 0.41, p = 0.50; task type: F(1,16) = 1.6, p = 0.18; TBS site X TBS scan: F(2,16) = 0.52, p = 0.60; TBS site X task type: F(2,16) = 1.7, p = 0.17; TBS scan X task type: F(1,16) = 2.16, p = 0.14; TBS site X TBS scan X task type: F(2,16) = 0.04, p = 0.96.

In summary, the analyses on the behavioral performance measures revealed: 1) a small but significant improvement in both accuracy and RT between the pre- and post-TBS scans, irrespective of TBS site and task type, which likely reflects a small training effect; and 2) TBS did not affect behavioral performance irrespective of stimulation site and/or task type. This may indicate that TBS caused changes in the magnitude of brain activity and the degree of functional connectivity that are not behaviorally relevant to configural or featural face processing or that the small changes in fMRI activity caused by TBS (0.17% signal change) were insufficient to produce differences in performance that could be detectible by the statistical analyses we performed. To distinguish between these two possibilities, we performed the following analysis within the face-selective ROIs of the right ventral stream (right aIT, FFA and OFA; defined during the localizer session). First, we extracted beta weight coefficients (% signal change) corresponding to the average activity for configural and featural face difference detections, separately for each participant. These beta weight coefficients were extracted from the pre-TBS scans of all sessions (left, right PPC and vertex) and were averaged (separately for each participant) to create an overall measure of fMRI activity corresponding to face processing within the face-selective regions of the right VOTC. We then conducted a linear regression analysis between these average measures of face-related activity in the ventral stream and RT performance (in msec) across participants. We used RT for the regression analysis as it had greater variability across participants compared to accuracy. We used a log10 transformation for the RT measures under the assumption that relatively large changes in RT are usually associated with small changes in BOLD magnitude. Unsurprisingly, the regression analysis was significant (r = 0.62, p = 0.02), indicating that activity within the face-selective regions of the right ventral stream predicted the participants’ behavioral performance: participants who were slower to respond to the face tasks had greater activity in response to face difference detections. The gradient of the regression line was 13.8 msec (1.14 at log10 converted to msec), indicating that, for this group of participants, for every one percent signal change in BOLD activity, there was a 13.8 msec change in RT. In our study, TBS caused a 0.17 percent signal change, which corresponds to an RT difference of only 2.35 msec. Thus, our findings are consistent with the assumption that the small changes in fMRI activity caused by TBS (0.17% signal change) were insufficient to produce differences in behavioral performance that could be detectible by the statistical analyses we performed.

### Analysis of eye fixations

Eye fixations were recorded, during the training session, immediately after participants completed their training on the configural and featural face processing tasks (details in Methods) because the MRI scanner available to us was not equipped with an eye-tracker. In the analyses, the number of fixations and fixation durations were the dependent measures with area of interest (AOI; see Methods) and task type (configural or featural face difference detections) as factors. All analyses were performed using linear mixed-effects models with participants added as a random variable. The analysis on the number of fixations yielded a significant main effect of AOI (F(63, 33643) = 444, p < 0.001) but not of task type (F(1, 33643) = 0.5, p = 0.39; on average, there were three fixations per trial for both face tasks), and there was no significant interaction between the factors (F(63, 33643) = 0.68, p= 0.97). The main effect of AOI showed that participants, irrespective of the type of face difference, mainly fixated the AOIs corresponding to the position of the eyes and the area around the nose of a face.

The analysis on the fixation duration data yielded similar results, with a significant main effect of AOI (F(63, 33643) = 424.2, p < 0.001) but not of type of face difference (F(1, 33643) = 0.01, p = 0.91; 316 msec per fixation for configural face difference detection trials and 315 msec per fixation for featural face difference detection trials), and there was no significant interaction between the factors (F(63, 33643) = 0.65, p= 0.99). The main effect of AOI indicated that fixations with the longest durations also occurred at the AOIs corresponding to the eye area of a face and the area around the nose.

In summary, participants focused their gaze on the same within-face locations for the same amount of time, irrespective of the face task.

## Discussion

Configural face processing is considered fundamental for face recognition and identification (Yovel and Kanwisher 2004; Maurer et al. 2007; Pitcher et al. 2007; Liu et al. 2010; Zhang et al. 2012, 2015). By using TBS in combination with fMRI, we investigated whether face-selective regions of the ventral visual pathway receive visuospatial information from location-processing regions of the dorsal visual pathway. Specifically, we explored whether TBS-induced interference of localized spatial-processing regions of the dorsal visual pathway affect the magnitude of fMRI activity and/or the degree of functional connectivity of localized face-processing regions of the ventral visual pathway during performance of configural and featural face difference detection tasks similar to those used in previous studies (Yovel and Kanwisher 2004; Duchaine et al. 2006; Yovel and Duchaine 2006; Maurer et al. 2007; Pitcher et al. 2007; Barton 2008; Liu et al. 2010; Renzi et al. 2013).

We found that TBS of spatial-processing regions of the left PPC increased activity in bilateral PPC and reduced activity in right aIT and right OFA for configural but not featural face difference detections. This differential effect of TBS between configural and featural face difference detections was specific to these face-selective ROIs in right VOTC. The pre- and post-TBS activity for configural and featural difference detections was comparable in all other ventral stream ROIs (right FFA, left aIT, FFA, and OFA), as well as in bilateral PPA in parahippocampal cortex.

These TBS effects on the magnitude of fMRI activity were bolstered by our findings on task-based functional connectivity: TBS of either the left or right PPC, but not of the vertex control site, reduced the level of functional connectivity between the face-selective regions in VOTC and the location-processing regions in PPC, but only during configural face difference detections. Functional connectivity was reduced between the right aIT (as the seed) and bilateral FFA, bilateral OFA, as well as bilateral PPC (the target sites). This reduced functional connectivity was significantly greater for configural than featural face difference detections.

In sum, our findings indicated that TBS-induced interference of location-processing regions of the dorsal stream selectively affected configural face processing in face-selective regions of the ventral stream. Consequently, the face-selective areas of the ventral stream and the location-processing areas of the dorsal stream must interact during configural face processing.

Our current findings extend our previous work (Zachariou et al. 2016), which showed that: 1) dorsal stream spatial-processing regions are more strongly activated by configural face processing than processing the shape features of a face; 2) brain activity evoked by configural face processing within these dorsal stream regions predicted the participants’ behavioral performance on the configural face task; and 3) rTMS delivered to the PPC (but not to the vertex control site) selectively impaired the participants’ performance during configural face processing but not during the processing of the shape of facial features.

Additionally, our current findings complement recent reports that the dorsal stream mediates object perception (Konen and Kastner 2008; Zachariou et al. 2014; Freud et al. 2016; Bracci et al. 2017; for a review, see Freud et al. 2016). Our findings are particularly consistent with a recent study by Van Dromme et al. (2016) which demonstrated that muscimol-driven inactivation of the macaque caudal intraparietal area decreases fMRI activation evoked by threedimensional object perception in anterior inferior temporal cortical regions. This raises the possibility that the dorsal stream contributes to object perception via the configural processing of an object’s shape features. Whether or not our findings extend to non-face object categories, however, is still an open question to be addressed in future studies.

### Limitations and caveats

Collectively, our findings indicated that, along with the contributions of the ventral visual pathway, the dorsal visual pathway also contributes to certain aspects of face perception. Three unexpected findings, however, remain: 1) TBS of the right PPC did not affect the magnitude of fMRI activity; 2) TBS did not affect behavioral performance on either face task; 3) TBS on the left PPC increased fMRI activity in bilateral PPC but decreased activity in face-selective areas of the right OFA and aIT during configural face difference detections.

We think that the first two of these issues may be explained by limitations of TBS: unlike face (and non-face) processing within the ventral stream, spatial processing within the dorsal stream is not characterized by well-defined functionally specialized brain regions (e.g., OFA and LOC) that can be directly interfered with by TBS (Ungerleider and Mishkin, 1982; Haxby et al. 1991; Cohen et al. 1996; Richter et al. 1997, 2000; Vingerhoets et al. 2002). On the contrary, based on our analysis, activity within the right PPC in response to both spatial judgments (localizer scans) and configural face processing (Pre-TBS scans) was extensive and covered multiple brain regions. It is challenging for TBS delivered to a single site to modulate activity over such a large region, with an area of stimulation of less than 10 mm^2^. In contrast, the activated region for both spatial judgments and configural face processing was much smaller in left PPC and, as such, TBS interfered with a larger portion of the configural face processing activity in left PPC compared to the right PPC.

Regarding the absence of a behavioral effect on either face task, given the extensive training our participants received, very large changes in fMRI activity (beyond what can be safely achieved with TBS) were needed to observe any differences in behavioral performance. Perhaps the use of rTMS, delivered safely at higher power output inside the MRI scanner using MRI-compatible TMS coils, would overcome these first two issues.

The third issue raised is more of a puzzle. Why did TBS on left PPC increase fMRI activity in bilateral PPC but decrease activity and functional connectivity in right VOTC areas? Because the TBS interfered with only a fraction of the overall PPC activity, it is possible that activity in PPC increased during configural face processing following TBS as a compensatory mechanism in response to the stimulation. That is, regions in left or right PPC unaffected by the TBS may have increased in activity to overcome the induced interference, enabling participants to perform the task. However, the mechanism by which this increased activity in PPC resulted in decreased activity in both magnitude and level of functional connectivity in right VOTC remains unknown, and implies that the interhemispheric network interactions between the dorsal and ventral streams is more complex than typically observed. It should be noted, however, that it is not uncommon for TMS to cause both increases and decreases in fMRI activity across neural systems, both within and across hemispheres. For instance, Plow et al. (2014) delivered rTMS (75% machine output at 1Hz for 15 minutes) on left IPS, immediately before participants performed a demanding visual attention task (visual motion tracking). This caused a local decrease in fMRI activity within the left IPS target site, but an increase in activity within the left superior parietal lobule and left medial precuneus, both of which are part of the same dorsal attention network (Corbetta and Shulman 2002; Corbetta et al. 2008). Importantly, however, activity in MT of the contralateral hemisphere, which is also involved in attentive motion tracking, decreased in activity. It is clear that the interpretation of such complex effects of TMS will ultimately depend on a deeper understanding of the underlying physiology.

### Patient populations

Lastly, our current findings have relevance to studies of face perception in patient populations, for example, patients with lesions of PPC. Based on our current findings, we would predict that these patients would exhibit face-processing impairments. The face-processing ability of patients with PPC lesions, however, is not routinely examined, presumably because the patients’ primary symptoms pertain to spatial, motor and/or attention-related deficits (Robertson et al. 1999; Goldenberg 2009; Kravitz et al. 2011). We think it would be valuable to re-examine patients with PPC lesions and evaluate their performance on face perception tasks.

## Acknowledgments

We wish to thank Dr. Maryam Vaziri-Pashkam for assistance with the TBS sessions and helpful comments on the manuscript. This work was supported by the NIMH Intramural Research Program (NIH Annual Report ZIA MH002918-09; NIH Clinical Study Protocols 93-M-0170 (NCT 00001360) and 12-M-0128 (NCT 01617408)).

